# Host-specific subtelomere: structural variation and horizontal transfer in asexual filamentous fungal pathogens

**DOI:** 10.1101/2023.02.05.527183

**Authors:** Xiaoqiu Huang

## Abstract

Several asexual filamentous fungal pathogens contain supernumerary chromosomes carrying secondary metabolite (SM) or pathogenicity genes as well as transposons. Supernumerary chromosomes have been shown in *in vitro* experiments to transfer from pathogenic isolates to non-pathogenic ones and between isolates whose fusion can result in vegetative or heterokaryon incompatibility (HET). However, much is still unknown about the extent of horizontal transfer of supernumerary chromosomes within and between asexual pathogenic populations in adaptation to their hosts. We investigated several asexual fungal pathogens for genomic elements involved in maintaining telomeres for supernumerary and core chromosomes during vegetative reproduction. We found that in vegetative populations or lineages with a nearly complete telomere-to-telomere genome assembly (e.g. *Fusarium equiseti* and five *formae speciales* of the *F. oxysporum* species complex), core and supernumerary chromosomes were flanked by highly similar subtelomeric sequences on the 3’ side and by their reverse complements on the 5’ side. This subtelomere sequence structure was specific to the host. We detected instances of recent horizontal transfer of regions of a supernumerary chromosome between distant populations in the *F. oxysporum* species complex, and we also found field isolates with two structurally different copies of a supernumerary chromosome in a young asexual population, raising the possibility that those copies originated from different lineages by intrastrain anastomosis. A large number of HET domain genes were located in SM/pathogenicity gene clusters, with a potential role in marking these gene clusters during vegetative reproduction. The emergence of novel asexual pathogenic populations by horizontal transfer of transposon-rich supernumerary chromosomes within and between populations poses challenges to the control and management of these pathogens.

## Introduction

Asexual filamentous fungal pathogens provide an intriguing system for studies of asexual pathogen population genomics through the characterization of genetic variation by genome sequencing. Asexual fungal pathogens, as well as sexual ones, possess plastic genomes with mosaic architecture allowing for their genes and other functional elements to evolve at various speeds (Croll & McDonald, 2012; Raffaele & Kamoun, 2012; Dong, Raffaele & Kamoun, 2015; Frantzeskakis, Kusch & Panstruga, 2019). Several asexual filamentous fungal pathogens contain supernumerary chromosomes (also known as accessory, conditionally-dispensable, or lineage-specific chromosomes), which are present in some but not all isolates (strains) of the species (Covert, 1998). Supernumerary chromosomes are enriched for transposons and pathogenicity genes that are present in some isolates but absent from closely related isolates of the same or different species (Ma et al., 2010; Rep & Kistler, 2010; Croll & McDonald, 2012; Raffaele & Kamoun, 2012; Huang et al., 2016; Vanheule et al., 2016; Vlaardingerbroek et al., 2016b). Pathogenicity gene clusters are also found in core chromosomes (often in AT-rich regions); they tend to be localized closer to the ends of core chromosomes (subtelomeres) (Ma et al., 2010; Wiemann et al., 2013; Dong, Raffaele & Kamoun, 2015; Niehaus et al., 2017). Supernumerary chromosomes have been shown in *in vitro* experiments to transfer between vegetatively incompatible isolates or to transfer from a pathogenic isolate to a non-pathogenic isolate (Horizontal Chromosome Transfer or HCT) in asexual filamentous fungi (He et al., 1998; Akagi et al., 2009; Ma et al., 2010; Vlaardingerbroek et al., 2016a; van Dam et al., 2017).

A rapid rate of supernumerary chromosomes loss during mitosis was observed in *F. oxysporum* (Vlaardingerbroek at al., 2016b) and in *Zymoseptoria tritici* (Möller et al., 2018). In *F. oxysporum*, phylogenetic studies suggest horizontal transfer of supernumerary chromosomes and supernumerary effector genes (van Dam et al., 2016; Fokkens et al., 2018), and supernumerary chromosomes are likely acquired by horizontal transfer through vegetative fusion of hyphae (Eschenbrenner et al., 2020). We hypothesized that supernumerary chromosomes were transferred as selfish genetic elements between isolates within an asexual population at a sufficiently high frequency to create structural variation within the population, because of a lack of the corresponding control present only in sexual reproduction. If the horizontal transfer of supernumerary chromosomes between isolates within asexual populations was prevalent or even continuous, we would expect to find, in a young asexual population with a low level of single nucleotide polymorphism (SNP) among its iso-lates, a high level of structural variation in supernumerary chromosomes between isolates in the population, two or more structurally different copies of a supernumerary chromosome within some isolates, and a high level of gene movement between supernumerary and core chromosomes within isolates, and a high level of structural variation between isolates in the population. To find instances of those events, we set out to explore the asexual species complex *Fusarium oxysporum* composed of many *formae speciales*, some of which contain polyphyletic lineages, and some of which contain monophyletic young lineages.

In the fission yeast *Schizosaccharomyces pombe*, adjacent to the end of the chromosome is the subtelomere containing species-specific homologous DNA sequences, whose functions include gene expression and chromosome homeostasis (Tashiro et al., 2017). In the budding yeast *Saccharomyces cerevisiae* and several filamentous fungi, the subtelomeres contain a helicase-like gene (Louis, 1995; Sánchez-Alonso & Guzman, 1998; Gao et al., 2002; Inglis et al., 2005; Rehmeyer et al., 2006). A DNA helicase regulates mutually exclusive expression of virulence genes in the subtelomeres of *Plasmodium falciparum* via heterochromatin alteration (Li et al., 2019). Because subtelomeres contain repetitive DNA, their DNA sequences are difficult to reconstruct by genome assembly programs, so they are sparsely represented in databases of whole genome sequence data (Wu et al., 2009). Only in *Colletotrichum higginsianum* isolate IMI 349063, was it reported that the supernumerary and core chromosomes possessed highly similar subtelomeric sequences containing a helicase-like gene (Dallery et al., 2017). This finding motivated us to investigate whether this subtelomeric structure is preserved within the same population or *forma specialis* so as to form its own unique type of subtelomeric sequences, and how those sequences arise.

Filamentous fungal subtelomeres contain repetitive DNA, which may be controlled by several genome defense mechanisms including Repeat-Induced Point mutation (RIP) (Cambareri et al., 1989; Gladyshev, 2017). Because RIP operates during the sexual cycle in filamentous ascomycetes, fungi currently undergoing sexual reproduction are unlikely to possess intact subtelomere domains; for example, the presence of nucleotide changes indicative of RIP was shown on a family of highly similar telomere-associated helicases in the fungus *Nectria haematococca* Mating Population VI (MPVI) (Coleman et al., 2009), and no repetitive DNA or helicase was found near the ends of chromosomes in the sexual filamentous fungus *Neurospora crassa* (Wu et al., 2009). Thus, a relevant question is whether repetitive DNA in the subtelomeres of an asexual fungal pathogen (with a strong RIP system) was acquired during the asexual cycle or during the emergence of the asexual pathogen.

We attempted to collect information on pathogenicity gene clusters in vegetative reproduction by exploring their connection to the genetic basis of vegetatively growing cells (in filamentous fungi) to distinguish self from nonself (allorecognition). During the vegetative growth phase, allorecognition can result in vegetative or heterokaryon incompatibility (HET) following fusion of genetically different cells, which disrupts growth and causes cell death (Glass & Dementhon, 2006; Paoletti & Saupe, 2009). However, heterokaryon incompatibility is suppressed following conidial anastomosis tube (CAT) fusion between vegetatively incompatible strains of *C. lindemuthianum* (Ishikawa et al., 2012), suggesting that HCT may result from CAT fusion (Manners & He, 2011). Heterokaryon incompatibility involves a protein partner with a HET domain as a trigger of programmed cell death (Paoletti & Clavé, 2007). A large number of HET domain genes were found in the genomes of asexual filamentous fungal pathogens, for example, 231 and 324 in two isolates of *Pyrenochaeta lycopersici* (Dal Molin et al., 2018). HET domain genes possess some characteristics of pathogenicity islands: they are highly variable between closely related isolates from different incompatibility groups, but they display trans-species polymorphism, where a HET domain gene from an isolate is more similar to one from an isolate of a different species than to ones from isolates of the same species (Muirhead, Glass & Slatkin, 2002). HET domain genes in three filamentous fungal species also exhibit a trend toward clustering near the ends of chromosomes (Zhao et al., 2015). The origins and functions of many HET domain genes in filamentous fungi remain unknown (Paoletti & Clavé, 2007; Smith & Lafontaine, 2013). A vexing question is whether HET domain genes tend to be found in pathogenicity gene clusters.

Recently, long-read sequencing such as Single Molecule, Real-Time (SMRT) Sequencing has resulted in genome assemblies that span repetitive elements and complex regions of lengths up to 30 kb. In this paper, we address the above questions by analyzing SMRT-based genome sequences of asexual fungal pathogens. We have made attempts to collect evidence to answer or shed light on these questions.

## Methods

Four large files of 14 to 33 GB are made available on Zenodo for reproducing many of the results in the manuscript. The file subtelomere.tar (Huang 2023a) should be unpacked on a Linux system, where the following data analysis programs need to be made available with the ‘module load’ command: Perl, Bowtie2, Samtools, GATK, Picard, Bedtools2, and Minimap2. After unpacking it, go to the directory subtelomere, which contains a number of subdirectories. One subdirectory is named data: the subdirectory subtelomere/data/reads/ is used to hold all compressed files of short reads in fastq format from the other three files (see below), and the subdirectory subtelomere/data/assembly/ includes the genome assemblies of the following isolates: *C. higginsianum* isolate IMI 349063, *F. equiseti* (Fe) isolate D25-1, *F. oxysporum* f.sp. *cepae* (Foc) isolate Fus2, *F. oxysporum* f.sp. *cubense* (Focb) race 1 isolate 160527, *F. oxysporum* f.sp. *cubense* tropical race 4 (TR4) isolate UK0001, *F. oxysporum* f.sp. *lycopersici* (Fol) race 3 isolate D11, *F. oxysporum* f.sp. *melonis* (Fom) isolate Fom001, *F. oxysporum* f.sp. radicis-cucumerinum (Forc) isolate Forc016 (see the file README in the subdirectory for more information).

The other subdirectories in the directory subtelomere contain instructions and scripts for reproducing many of the results in the manuscript. The subdirectory subtelomere/TwoCopies/ contains two subdirectories of instructions and scripts for reproducing the results in Table 3 and Figure 2. The subdirectory subtelomere/TEs/ contains four subdirectories, each of which provides instructions and scripts for estimating the copy numbers of one or two transposons in Table 6. The subdirectory subtelomere/SVs/ explains how the results in Table 5 could be reproduced. The subdirectory subtelomere/Fol/ contains three subdirectories, each of which includes information for reproducing one of the three columns in Table 2. The subdirectory subtelomere/Forc/ demonstrates how some of the programs and scripts developed by the author are used to analyze a genome assembly of Forc isolate Forc016. The subdirectory subtelomere/pub/ contains the source and executable code of those programs. See the README and z.cmd files in each leaf subdirectory for more information.

Each of the other three files is packed from a directory of pairs of compressed files in fastq format containing paired-end reads. After unpacking each file, all files in the directory need to be moved to the subdirectory subtelomere/data/reads/. The file Data.One.Focb.tar (Huang 2023b) contains 16 files of paired-end reads Focb TR4 isolates II-5, S1B8, JV14, FOC.TR4-5, FOC.TR4-1, Col2, Col4, Col17. The file Data.One.Focb-2.tar (Huang 2023c) contains 22 files of paired-end reads from Focb race 1 isolate N2, and Focb TR4 isolates Hainan.B2, My-1, La-2, Vn-2, Leb1.2C, JV11, Phi2.6C, Pak1.1A, UK0001. The file reads.tar (Huang 2023d) contains 44 files of paired-end reads from Fol isolates Fol069, Fol072, Fol4287, and Forc isolates Forc016, Forc024, Forc031.

We also obtained the datasets of short reads for the following isolates (by their SRA accessions) from Sequence Read Archive (SRA) at NCBI: *C. higginsianum* isolate MAFF 305635 (SRR6412364); *F. equiseti* isolate CS5819 (SRR5194938); *F. oxysporum* isolate Fo47 (SRR306682, SRR306671, SRR306668, SRR306667, SRR306657); *F. oxysporum* f.sp. *cepae* isolates 125 (SRR4408417), A23 (SRR4408418), Fus2 (SRR4408416); *F. oxysporum* f.sp. *lycopersici* isolates Fol007 (SRR3142257, SRR3142258), Fol014 (SRR307256, SRR307271), Fol026 (SRR307236, SRR307255, SRR307293); Forc016 *in planta* RNA-seq (SRR5666309), *F. oxysporum* f.sp. *melonis* isolates Fom004 (SRR1343458), Fom005 (SRR1343500), Fom006 (SRR1343569), Fom012 (SRR1343574), Fom013 (SRR1343575), Fom016 (SRR1343577). Isolate Fo47 is non-pathogenic, the pathogenicity status of isolate CS5819 is unknown, and all the other isolates are pathogenic.

Pairwise comparisons of genome assemblies were performed with MUMmer3 (Kurtz et al., 2004), DDS2 (Huang et al., 2004) and Minimap2 (Li, 2018). Comparisons of a genome assembly with a database of genome assemblies were made with Blastn (Altschul et al., 1990). An alignment of two genomic regions with sections of similar regions separated by sections of different regions was computed with GAP3 (Huang & Chao, 2003). Multiple local alignments between two sequences were computed with SIM (Huang & Miller, 1991). Genes in a genome assembly were identified with Augustus (Stanke & Waack, 2003) and AAT (Huang et al., 1997). Domains in protein sequences were found with HMMER (Finn, Clements & Eddy 2011).

A chromosome or contig is said to terminate at its 3’ (or 5’) end in a telomeric repeat if a 3’ (or 5’) terminal region of 240 bp contains at least 16 occurrences of the word TTAGGG (or CCCTAA). The subtelomere of a chromosome or contig is defined to be a longest terminal region that ends in a telomeric repeat and is highly similar to terminal regions elsewhere.

Short reads were mapped onto a genome assembly as a reference, as previously described (Huang, 2014; Huang et al., 2016). Note that the highly similar subtelomeres of core and supernumerary chromosomes in the genome assembly were not covered by any short reads, because no short reads can be uniquely mapped to any duplicated regions in the reference. To determine whether a subtelomere was present in the genome represented by short reads, the sequence of the subtelomere was used alone as a reference for mapping the short reads. To estimate the copy number of the subtelomere in the genome represented by short reads, the average coverage depth of the subtelomere by short reads was calculated, and also calculated (for comparison) was that of a genomic region containing the *EF-1α* gene. The estimated copy number of the subtelomere was defined as the average coverage depth for the subtelomere divided by that for the *EF-1α* gene region. Note that the coverage depth at each base position of a genomic region was calculated by using Bedtools with the genomecov subcommand (Quinlan & Hall, 2010). The copy number of a transposon in the genome was estimated in a similar way to that of a subtelomere.

We also used an alternative approach to calling sequence variants between the reference genome and the genome of another isolate represented by short reads. The only difference between the previous and alternative approaches was that the alternative approach used GATK (Van der Auwera et al., 2013) with the HaplotypeCaller option to call sequence variants including SNPs. The alternative approach was used in all variant analysis tasks involving Focb race 1 or TR4 isolates.

## Results

### Similarities between subtelomeres within *formae speciales*

We studied similarities between subtelomeres within *F. oxysporum* f.sp. *radicis-cucumerinum* (Forc). We selected three Forc isolates: Forc016, Forc024 and Forc031 (van Dam et al., 2016). A genome assembly of Forc016 consisted of 11 core chromosomes, one supernumerary chromosome (chromosome RC), and a number of unplaced contigs (van Dam et al., 2017). Chromosome RC, assembled from telomere to telomere, was flanked on either side by 13-kb reverse complementary sequences at 99.96% complementarity (Fig. 1A). Of the core chromosomes, 7 were assembled from telomere to telomere and flanked on either side by 13-kb reverse complementary sequences at 99.98-100% identity to and in the same orientation as those of chromosome RC (Fig. 1B), and the other 4 were flanked on one side by subtelomeres of this common type. Seven of the unplaced contigs were flanked on one side by common subtelomeres at 99.68-100% identity. The common Forc016 subtelomere was also present in multiple copies in the other two isolates; its estimated copy number was 26 in isolates Forc024 and Forc031, and 29 in isolate Forc016.

**Figure 1.**
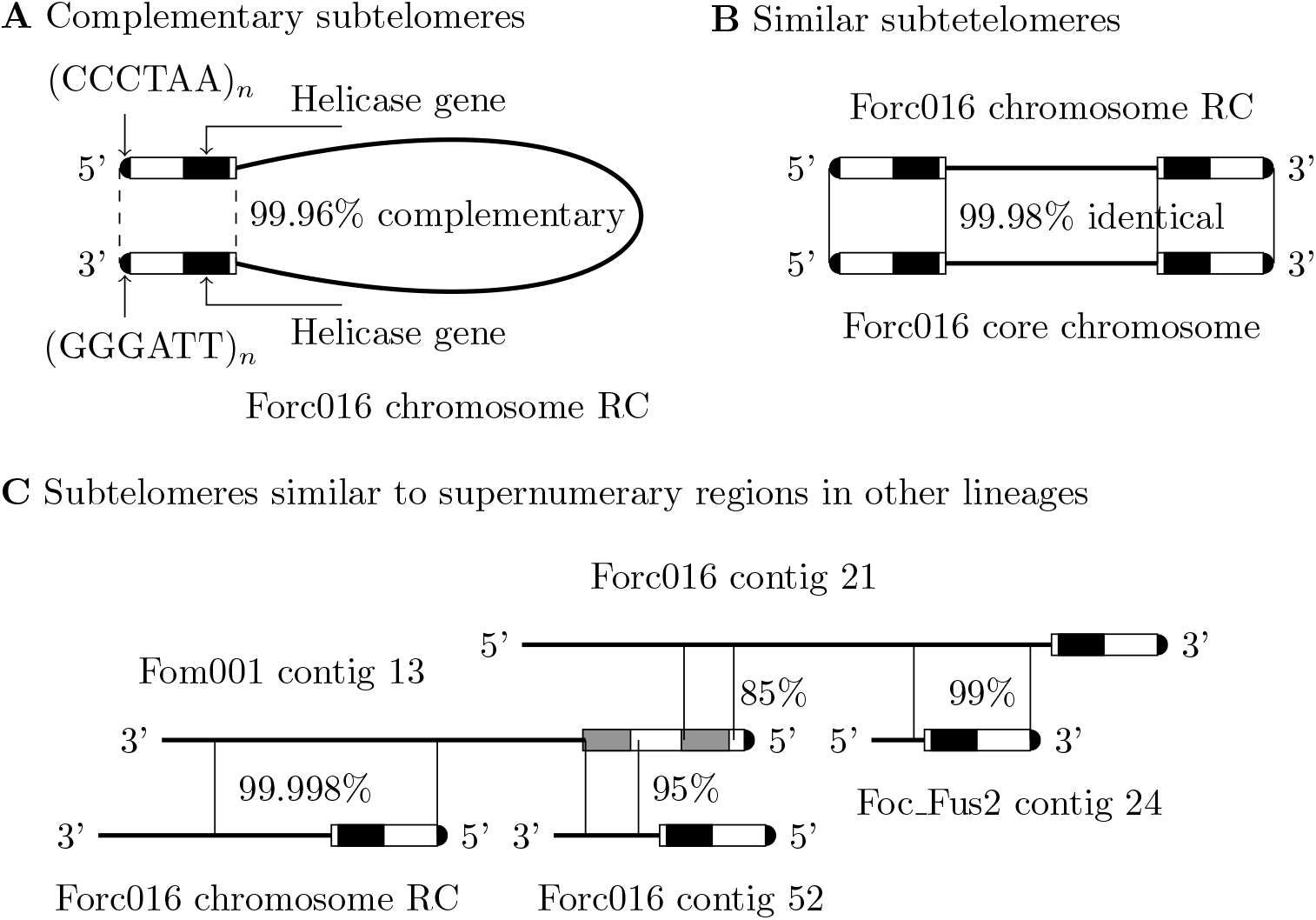
Three types of subtelomere sequence similarities. Each chromosome or contig is represented by a thick solid line with its name placed above or below the line. Similar regions between chromosomes/contigs are indicated by two vertical thin solid lines at their boundaries, with the percent identity given next to the lines. The subtelomere ends in a telomeric repeat (indicated by a half-filled circle) and contains a helicase gene (shown by a black rectangle). The figure is not drawn to scale. (*A*) The 5’ and 3’ subtelomeres of one single strand of Forc016 chromosome RC are complementary (indicated by two dashed lines at their boundaries) when read from both ends, making it possible for the supernumerary chromosome to repair its subtelomeres by forming a subtelomere hairpin if one of them is shortened. (*B*) The subtelomeres of core and supernumerary chromosomes at the same end are nearly identical, making it possible for them to exchange genes and repair subtelomeres by homologous recombination. (*C*) A subtelomere (in multiple copies) from one vegetative lineage is homologous to a non-terminal region (in a single copy) of a supernumerary chromosome or contig from another lineage. The subtelomere of Fom001 contig 13 contains a duplication (shown by a pair of gray rectangles), instead of a helicase gene.

The comparison of the common Forc016 subtelomere with the genome assemblies of all isolates in the *Fusarium* genus revealed that the subtelomere was 100% identical over 12.9 kb to 6 *F. oxysporum* f.sp. melonis (Fom) isolates. Its estimated copy number was 2 or 3 in isolates Fom005, Fom006, Fom012, Fom013, and Fom016. Chromosome RC of isolate Forc016 was much more conserved than its core chromosomes between Forc016 and these Fom isolates. For example, there were no SNPs between Forc016 and each of these isolates over a 31-kb region (starting at 766.8 kb) of Forc016 chromosome RC. In contrast, there were at least 60 SNPs over a 20.6-kb region (containing the conserved *EF-1α* gene) between Forc016 and each of these isolates. Note that both Forc and Fom isolates caused severe disease in musk melon (van Dam et al., 2016), a common host for both *formae speciales*. The Forc016 subtelomere was 95-96% identical over 12.4 kb to 4 isolates from other *formae speciales*; it had no strong matches to other isolates.

In isolate Forc016, some chromosomes and contigs were also homologous over a length of 4 kb beyond the 13-kb subtelomere, with numerous G-C to A-T nucleotide changes indicative of possible RIP. For example, all next to their 3’ 13-kb subtelomeres, a region of chromosome 4 was 90% identical to one of chromosome 10 with 394 G-C to A-T nucleotide changes out of a total of 396 substitutions, 91% identical to one of contig 7 with 329 out of 332, and 88% identical to one of contig 3 with 453 out of 456. An explanation for this observation is that RIP may have operated on the parts of these subtelomeres in an ancestral state.

Five of the unplaced contigs were not present in isolates Forc024 and Fore031. Two of these 5 contigs end in a telomeric repeat: the 3’ end of contig 55 and the 5’ end of contig 52. The 3’ subtelomere of contig 55 in forward orientation was 100% identical over 7,100 bp to the 5’ one of contig 52 in reverse orientation, and the subtelomere contained a gene encoding for a protein of 1,531 residues with Helicase C (e-value = 1.2e-09), DEAD (e-value = 3.2e-09) and ResIII (e-value = 5.3e-05) domains. This new subtelomere was not similar to those of the core chromosomes. The longest of the 5 contigs, contig 53 of 987.8 kb, was 95-98% identical over 36% of its length to supernumerary Fol D11 contig 38, and 93-99% identical over 21% of its length to supernumerary Fol D11 chromosome 14. Thus, isolate Forc016 appeared to contain a supernumerary chromosome with a different host-specific subtelomere, which might have been acquired by horizontal transfer and maintained with the palindrome subtelomere at its ends.

Subtelomere similarities of the type in Fig. 1B within Forc, Fol, and other *F. oxysporum formae speciales*, as well as within other asexual populations, are summarized in Table 1. Refer to Methods for abbreviations for the names of all *F. oxysporum formae speciales* and additional information on all the isolates. In isolate Forc016 in Table 1, its entire reference subtelomere was present in multiple copies in the other two isolates. In isolate Focb TR4 UK001, part of its reference subtelomere at 4000-9000 bp was covered by all of the other 15 isolates, but part of it at 1-3000 bp was not covered by 7 of the 15 isolates, and most of it was not covered by reads from isolate Pak1.1A, which was not among the 15 isolates.

**Table 1.** Subtelomere similarities in *F. oxysporum formae speciales* and other asexual populations

| Isolate | Reference<br>subtelomere | No. in<br>this <sup>a</sup> | Subtelomere<br>similarities <sup>b</sup> | No. in<br>others <sup>c</sup> |
| --- | --- | --- | --- | --- |
| Fol D11 | 3' chromosome 1 | 22 | 99.89-99.98% | 25-35 (6) |
| Forc016 | 3' chromosome RC | 26 | 99.68-100% | 26 (2) |
| Fom001 | 5' contig 13 | 17 | 98.69-99.76% | 24-44 (6) |
| Fom001 | 5' contig 22 | 6 | 99.27-99.96% | 4-7 (5) |
| Focb race 1 160527 | 3' contig 2 | 20 | 98.83-99.98% | 26 (1) |
| Focb TR4 UK0001 | 5' contig 13 | 22 | 99.47-100% | 22-30 (15) |
| Foc Fus2 | 3' contig 24 | 4 | 98.89-99.97% | 25-30 (3) |
| Fe D25-1 | 3' scaffold 15 | 4 | 99.63-99.73% | ? (0) |
| IMI 349063 | 3' chromosome 12 | 8 | 99.32-99.94% | 22 (1) |
| IMI 349063 | 3' chromosome 11 | 3 | 99.87-99.91% | 20 (1) |

<sup>a</sup> The number of subtelomeres other than the reference subtelomere in the genome assembly of this isolate.
<sup>b</sup> The range of percent identities between the reference subtelomere and each of the other subtelomeres in the genome assembly of this isolate.
<sup>c</sup> The range of estimated subtelomere numbers in the genome datasets of other isolates (with the number of isolates given in parentheses) in the *F. oxysporum forma specialis* (or species) of the isolate named in column 1. In some cases (e.g., Focb race 1 160527, Focb TR4 UK0001, IMI 349063), only part of the reference subtelomere was present in multiple copies in the isolates (see details in text).

To see how unique the 3’ subtelomere of D11 chromosome 1 was to Fol, its sequence was compared to all genome assemblies (over 200) in the *Fusarium* genus. Of the top 35 matches, 19 were to the genome assembly of isolate D11, which were described above, and the 16 were 99.66-99.99% identical over a length of 7.4-10.7 kb to genomic regions from other isolates in this *forma specialis*. The next 6 best matches were 97-98% identical over 6.5-8.3 kb to isolates in other *formae speciales* of *F. oxysporum*.

### Some supernumerary regions much more conserved than all core regions

First we examined isolates D11, Fol4287, Fol069 and Fol072 in *forma specialis* Fol. We checked whether supernumerary D11 chromosome 14 and contig 38 were present and conserved in isolates Fol069 and Fol072, which were not in the same lineage as isolates D11 and Fol4287. For each of isolates Fol4287, Fol069 and Fol072, short reads from the isolate were mapped onto the Fol D11 genome assembly (Henry et al., 2019) as a reference, the mapped reference was partitioned into disjoint windows of maximum lengths such that each position in the window was covered at a depth greater than or equal to *adc/wca*, and each window of at least length *wlen* was selected to calculate its SNP rate, where *adc* is the average depth of coverage over the reference genome for the isolate, and *wlen* and *wca* are parameters. Note that windows were introduced to select regions of the reference that were unique and less variable between Fol D11 and the other isolates. The parameters *wlen* and *wca* were set to 10,000 bp and 5, respectively, to ensure that there were a sufficient number of windows for each isolate (Table 2). Supernumerary D11 chromosome 14 and contig 38 were present in each isolate, based on their percent coverage by short reads from the isolate (Table 2), which was the percentage of the chromosome or contig covered at a depth greater than or equal to *adc/wca*. They were more conserved than the core chromosomes between Fol D11 and each of isolates Fol069 and Fol072, by the weighted mean of the SNP rates of windows in each of them in comparison to that in the genome (Table 2), where the weight for each rate was the size of its window divided by the sum of all window sizes. This was unexpected, as supernumerary chromosomes were much more variable than core chromosomes among isolates from the same species (Huang et al., 2016; Vlaardingerbroek et al., 2016b).

**Table 2.** Mean SNP rates between Fol D11 and each of Fol4287, Fol069 and Fol072

| Coverage and SNP distribution | Fol4287 | Fol069 | Fol072 |
| --- | --- | --- | --- |
| Average depth of coverage (*adc*) | 90.6 | 64.7 | 57.6 |
| Percent coverage of D11 chr 14<sup>a</sup> | 64% | 52% | 51% |
| Percent coverage of D11 ctg 38 | 68% | 66% | 39% |
| Nos. of windows in the genome/chr 14/ctg 38 | 645/34/24 | 252/16/23 | 225/16/16 |
| Mean SNP rate of windows in the genome | 0.00001 | 0.00349 | 0.00361 |
| Mean SNP rate of windows in chr 14 | 0.00009 | 0.00001 | 0.00001 |
| Mean SNP rate of windows in ctg 38 | 0.00003 | 0.00003 | 0.00007 |

<sup>a</sup> The percent coverage of D11 chromosome (chr) 14 by short reads from the isolate was the percentage of the chromosome covered at a depth greater than or equal to *adc*/5.

**Figure 2.**
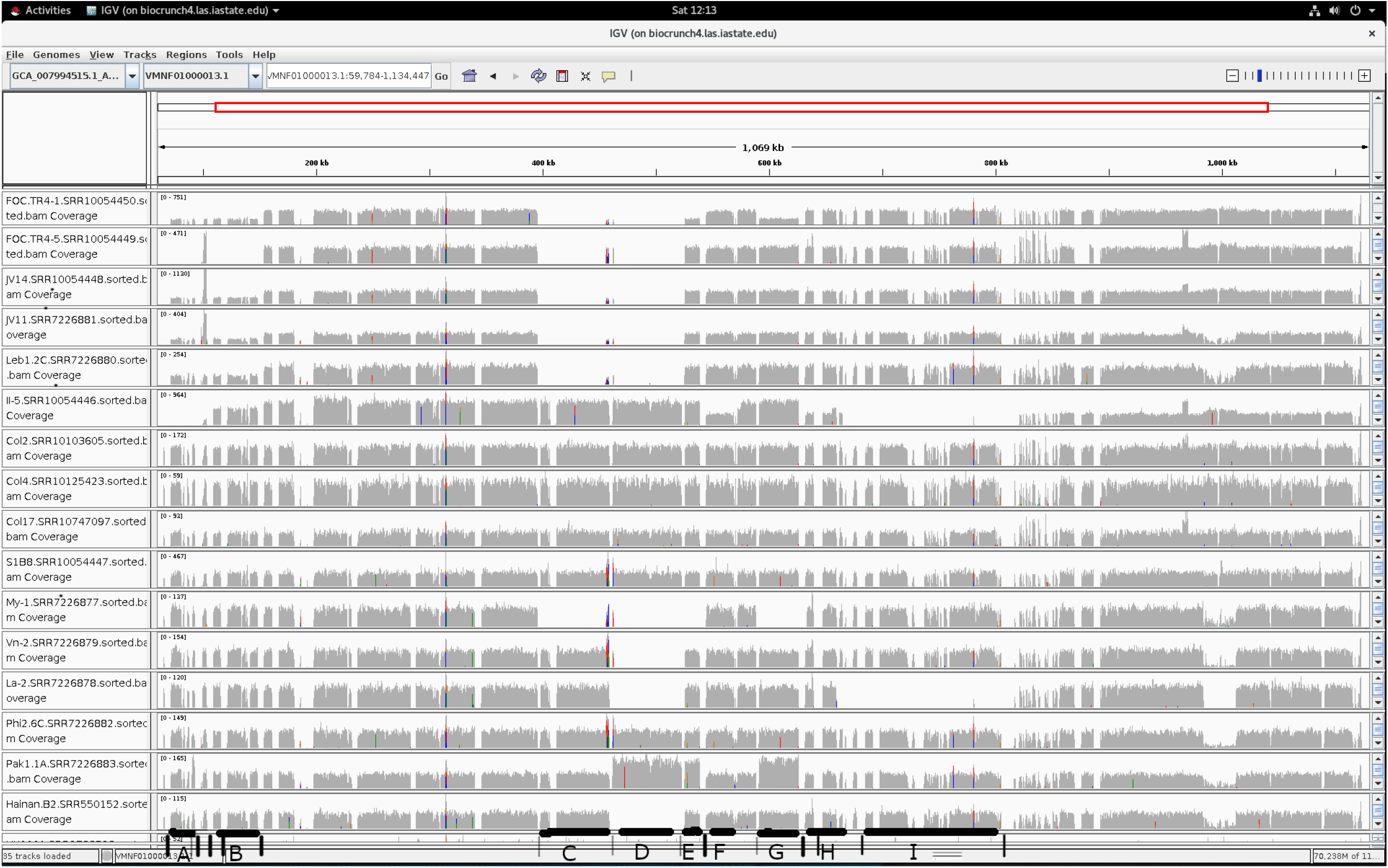
Coverage of a large portion of supernumerary TR4 UK0001 contig 13 (shown in the top horizontal panel) by short reads from 16 TR4 isolates, with each isolate in a separate horizontal panel. Each left panel shows a prefix of an alignment file name, made up of of the name and NCBI SRA accession number of the isolate. Nine regions of contig 13 in which CNVs were observed among the isolates are marked with thick horizontal lines and designated with the letters A through I. These nine regions were located next to copies (with 0 to 3 nucleotide differences) of three repetitive elements of 3482, 5682 and 6122 bp, each containing a transposon, with the copies of the 3482-bp element indicated by thin vertical lines at the bottom, those of the 5682-bp element by medium vertical lines, and those of the 6122-bp element by thick vertical lines.

Next we considered race 1 and TR4 isolates in *forma specialis* Focb. We investigated whether regions from supernumerary race 1 160527 contig 2 or TR4 UK0001 contig 13 were more conserved than ones from the other contigs, between the race 1 160527 and TR4 UK0001 assemblies. We compared the two genome assemblies and found 105 matches of at least length 10 kb with no insertions/deletions, and partitioned them into two groups: group 1 of 48 matches with one or two regions of the two supernumerary contigs, and group 2 of 57 matches with no regions from those two contigs. The match with the lowest SNP rate in group 2 was of length 10,360 bp at a rate of 0.0032. Group 1 had 21 perfect matches with a SNP rate of 0, and all but one (at length 10,597 bp) of them contained both regions of the two supernumerary contigs. The longest of those matches was 60,915 bp, and the next four perfect ones were 57,966, 42,438, 36,352 and 33,736 bp in length. The SNP rate of the longest perfect match in group 1 was at least 193 times lower that that of the best match in group 2. In addition, by mapping short reads from TR4 UK0001 onto the race 1 160527 assembly as a reference, we obtained a SNP rate of 0.013950 (530, 040*/*37, 995, 238) across the genome between race 1 160527 and TR4 UK0001, with the SNP rate being defined as the number of high-quality SNPs divided by the number of sufficiently covered positions in the genome. Thus, some regions of the two supernumerary contigs were much more conserved than all regions of the core chromosomes between Focb race 1 160527 and TR4 UK0001, suggesting that those regions were acquired by at least one of the two lineages through horizontal transfer. These two lineages are shown below to have separated from each other early in the F. oxysporum species complex.

To see where race 1 160527 and TR4 UK0001 were located phylogenetically along with other F. oxysporum isolates, we utilized phylogenetic trees of 99 F. oxysporum isolates constructed using three different types of data: the whole genome, the conserved region of the mitochondrial genome and eight informative nuclear genes (Achari et al., 2020). Most of the 99 isolates were classified into three clades named Clades 1-3 that were concordant in all the phylogenetic trees, where Clade 1 had diverged early from Clades 2 and 3, which have separated from each other more recently (Achari et al., 2020). TR4 UK0001 was phylogenetically placed in Clade 1, because Clade 1 contained two isolates (F oxy NE 5689 and F oxy CAN 42420) such that one of them was closer to TR4 UK0001 than the other was. Similarly, race 1 160527 was phylogenetically placed in Clades 2 or 3, because Clades 2 and 3 contained two isolates (F oxy NE 6324 and F oxy CAN 43194) such that one of them was closer to race 1 160527 than the other was. Note that the evolutionary distances between the above isolates were estimated by computing the average percent identity of gap-free matches of at least 10,000 bp between the genome sequences of those isolates.

A low level of diversity among Focb TR4 isolates was observed with only 251 SNPs found across the genome (at a SNP rate of 0.000005) among 8 Focb TR4 isolates (Zheng et al., 2018). However, there was a much higher level of diversity between isolates in Focb race 1. Between isolate 160527 and another race 1 isolate N2 (Guo et al. 2014), we observed a SNP rate of 0.0059 over a 10-kb region containing the conserved *EF-1α* gene, a similar SNP rate over the 10-kb region (160527 contig 5 at 1,359,659-1,370,018 bp) in the match with the lowest SNP rate in group 2 (see above), and a SNP rate of 0.006343 across the whole genome. In comparison, between the two isolates, we observed a SNP rate of 0.000535 (819*/*1, 530, 705) over a 3.1-Mb 3’ segment of supernumerary race 1 160527 contig 2. An examination with the Integrative Genomics Viewer (IGV) confirmed that that all large segments of the other contigs in the 160527 assembly had a much higher SNP density than the 3’ segment of supernumerary contig 2. Similarly, we observed a SNP rate of 0.00012 over a 3’ 8.3-kb subtelomere of Focb 160527 supernumerary contig 2, after mapping short reads from isolate N2 onto a 3’ 85.8-kb end of this contig. Furthermore, a 5.310-bp region of this subtelomere located at 3.1 kb from the 3’ end of the contig, contained no SNPs, with its depth of short read coverage being 26 times the coverage depth of the *EF-1α* locus. Most significantly, a SNP rate of 0 was observed in a 50,826-bp region of supernumerary race 1 160527 contig 2 at 3,620,000-3,670,825 bp between race 1 160527 and N2, where this region was sufficiently covered by short reads from N2. Thus, some large regions of supernumerary contig 2 were much more conserved than all large regions of the core chromosomes between the two isolates in Focb race 1, which suggests that those regions were acquired by at least one of the two lineages through horizontal transfer. In particular, these observations suggest that the subtelomere was acquired by horizontal transfer and duplicated at the ends of many chromosomes in at least one of the two lineages.

### Two or more copies of a supernumerary chromosome in isolates

We found more than one copy of a supernumerary chromosome in several Focb TR4 isolates. As indicated above, Focb TR4 isolates had much lower genetic diversity than Focb race 1 isolates, suggesting that TR4 isolates emerged much more recently than race 1 isolates. By mapping short reads from Focb TR4 isolates onto a genome assembly of TR4 UK0001, we observed, in five of the TR4 isolates, significant variation in coverage depth between some of the nine regions of supernumerary TR4 UK0001 contig 13 (Fig. 2). We compared the average depth of coverage of TR4 UK0001 contig 13 with that of core TR4 UK0001 contig 6 for each TR4 isolate (Table 3); the results suggest that more than one copy of supernumerary contig 13 were present in four (FOC.TR4.1, JV14, JV11, Leb1.2C) of the five TR4 isolates and that more than two copies were present in the other isolate (II-5). The copies in each of these five isolates were different in some of the nine regions of UK0001 contig 13 (Table 3 and Fig. 2), making it possible for more variants to be generated by homologous recombination. Table 3 reports copy-number variations (CNVs) of contig 13 in the same isolate and in the same clade, and common CNVs of contig 13 in different clades, raising the question of whether differing copies of contig 13 were acquired by horizontal transfer.

**Table 3.** Average coverage of Focb TR4 UK0001 contig (ctg) 13 and ctg 6 by short reads from other Focb TR4 isolates

| Isolate<sup>a</sup> | Average coverage<br>for ctg 13 | Average coverage<br>for ctg 6 | Haplotype/genotype<br>of ctg 13<sup>b</sup> |
| --- | --- | --- | --- |
| FOC.TR4-1 | 305.5 | 155.9 | $A^{\pm}B^{\pm}C^{-}D^{-}E^{\pm}F^{+}G^{\pm}H^{+}I^{+}$ |
| FOC.TR4-5 | 201.1 | 177.7 | $A^{-}B^{-}C^{-}D^{-}E^{+}F^{+}G^{+}H^{+}I^{+}$ |
| JV14 | 355.9 | 172.1 | $A^{\pm}B^{\pm}C^{-}D^{-}E^{+}F^{+}G^{+}H^{+}I^{+}$ |
| JV11 | 105.2 | 49.6 | $A^{\pm}B^{\pm}C^{-}D^{-}E^{+}F^{+}G^{+}H^{+}I^{+}$ |
| Leb1.2C | 106.8 | 50.6 | $A^{\pm}B^{\pm}C^{-}D^{-}E^{+}F^{+}G^{+}H^{+}I^{+}$ |
| II-5 | 376.8 | 134.7 | $A^{-}B^{\pm}C^{+}D^{+}E^{+}F^{\pm}G^{+}H^{\pm}I^{-}$ |
| Col2 | 82.0 | 72.8 | $A^{+}B^{+}C^{+}D^{+}E^{+}F^{+}G^{+}H^{+}I^{+}$ |
| Col4 | 31.4 | 28.1 | $A^{+}B^{+}C^{+}D^{+}E^{+}F^{+}G^{+}H^{+}I^{+}$ |
| Col17 | 30.8 | 26.9 | $A^{+}B^{+}C^{+}D^{+}E^{+}F^{+}G^{+}H^{+}I^{+}$ |
| S1B8 | 151.5 | 142.4 | $A^{+}B^{+}C^{+}D^{+}E^{+}F^{+}G^{+}H^{+}I^{+}$ |
| My-1 | 64.1 | 57.9 | $A^{+}B^{+}C^{-}D^{-}E^{-}F^{+}G^{-}H^{+}I^{+}$ |
| Vn-2 | 62.9 | 58.4 | $A^{+}B^{+}C^{+}D^{-}E^{+}F^{+}G^{+}H^{+}I^{+}$ |
| La-2 | 63.7 | 57.2 | $A^{+}B^{+}C^{+}D^{-}E^{+}F^{+}G^{+}H^{+}I^{-}$ |
| Phi2.6C | 58.6 | 52.7 | $A^{+}B^{+}C^{+}D^{+}E^{+}F^{+}G^{+}H^{+}I^{+}$ |
| Pak1.1A | 61.5 | 51.8 | $A^{+}B^{+}C^{+}D^{+}E^{+}F^{+}G^{+}H^{+}I^{+}$ |
| Hainan.B2 | 41.8 | 37.8 | $A^{+}B^{+}C^{+}D^{-}E^{+}F^{+}G^{+}H^{+}I^{+}$ |

<sup>a</sup> Top 15 isolates, excluding the one in the last row, are arranged into three groups delimited by horizontal lines, with each group corresponding to one of the three separate clades in a phylogenetic tree of these isolates (Maymon et al., 2020).
<sup>b</sup> Nine regions of Focb TR4 UK0001 ctg 13 are marked with thick lines and designated with *A* through *I* (see Fig. 2). For any of the nine regions (say, region *A*) and an isolate, the notation $A^{+}$ denotes that region *A* is present in one or two copies of ctg 13 in the isolate, and $A^{-}$ denotes that region *A* is absent from one or two copies of ctg 13 in the isolate. The notation $A^{\pm}$ denotes that the isolate contains two copies of ctg 13, with region *A* present in one copy and absent from the other copy.

Figure 2 shows a few common recombination types of regions A through I, some of which were shared by isolates in different clades of a phylogenetic tree of 15 Focb TR4 isolates, with the isolates in each clade shown in Table 3. For example, the deletion of regions C and D was shared by isolate My-1 in one clade and by five isolates including FOC.TR4-1 in another clade; the deletion of regions E and G was shared by isolate My-1 in one clade and isolate FOC.TR4-1 in another clade; and the deletion of region I was shared by isolate La-2 in one clade and isolate I-5 in another clade. In addition, regions D and I contained a perfect match of 33,736 bp and a perfect match of 57,966 bp, respectively, between Focb TR4 UK0001 supernumerary chromosome 13 and Focb race 1 160527 supernumerary contig 2, in spite of a genome-wide SNP rate of 0.013950 between Focb TR4 UK0001 and race 1 160527 (see above). Both of these matches were mentioned above. Both matches were likely to be due to horizontal transfer between the two races, which were located in different clades of *F. oxysporum* isolates, as indicated above. Regions A through I were next to repetitive elements of Focb TR4 UK0001 supernumerary contig 13, shown by vertical bands of gaps across panels in Fig. 2, because they were not covered by any short reads. We found three types of repetitive elements each with five or more copies in contig 13: two of them were of 3482 and 5682 bp, each containing an LTR retrotransposon or part of it, and the other was of 6122 bp, containing a helitron. Moreover, identical or nearly identical copies of these three types of transposons were also found in core chromosomes in Focb TR4 UK0001, and in supernumerary contig 2 and core contigs in Focb race 1 160527, but not in other *formae speciales* of *F. oxysporum* (Table 4). Some of the copies in Focb TR4 were located at structural variants in core chromosomes among the 16 Focb TR4 isolates (Table 4).

**Table 4.** The numbers of copies identical and nearly identical to sample copies of three transposons in Focb TR4 UK0001 and race 1 160527 as well as the numbers of copies located at CNVs of lengths 15 to 170 kb among 16 TR4 isolates

| Transposon sample<br>in TR4 UK0001<sup>a</sup> | PCN/CN (SPCN/SCN)<br>{ICN} in TR4 UK0001<sup>b</sup> | CN (SCN) {ICN}<br>in race 1 160527 | CN in<br>others |
| --- | --- | --- | --- |
| ctg 13: 87,797 (3482 bp) | 5/9 (5/6) {9} | 34 (25) {29} | 0 |
| ctg 13: 809,230 (5682 bp) | 27/42 (3/5) {18} | 3 (3) {2} | 0 |
| ctg 13: 66,262 (6122 bp) | 13/32 (7/8) {8} | 18 (12) {0} | 0 |

<sup>a</sup> Shown are the start coordinate and length (in parentheses) of a sample copy for each of the three types of transposons in supernumerary TR4 UK0001 contig (ctg) 13.
<sup>b</sup> CN, the number of copies in TR4 UK0001 (or in race 1 160527 or in other non-Focb *F. oxysporum* isolates) that were identical (ICN) or nearly identical (with at most 5 nucleotide differences) to the sample copy; PCN ( ≤ CN), the number of such copies in TR4 UK0001 that were located at CNVs of lengths 15 to 170 kb among the 16 TR4 isolates. In TR4 UK0001, SPCN and SCN denote such PCN and CN numbers limited to ctg 13; in race 1 160527, SCN denotes such a CN number limited to ctg 2.

### Structural variants in supernumerary and core chromosomes

We found 118 SNPs between the UK0001 reference and 16 other TR4 isolates in the supernumerary chromosome. We also found more than 118 kb of nucleotides affected by presence/absence structural variants between the UK0001 reference and 16 TR4 isolates in the same chromosome (Fig. 2). Thus, the total number of nucleotides affected by structural variants was 1000 times more than the number of SNPs in the supernumerary chromosome. To compare the amount of structural variation to the number of SNPs in the core chromosomes, we selected 8 pairs of TR4 isolates, 6 of them in the same clade. For each pair of TR4 isolates, we found SNPs between the UK0001 reference and each isolate in the pair, and kept the SNPs unique to each isolate by removing the SNPs common to both isolates in the pair. The amount of structural variation between the isolates in each pair was estimated by finding presence/absence polymorphisms (PAPs) unique to each isolate. A region of the UK0001 reference was present in isolate A but absent from isolate B if every position in the region was covered by short reads from A to at least .95 times the average read depth for A, and by short reads from B to at most 0.01 times the average read depth for B. A region of at least 200 bp was a PAP unique to one isolate in the pair if it was present in the isolate but absent from the other in the pair and no larger region containing it can meet this requirement. The results for the 8 pairs of TR4 isolates are shown in Table 5. Thus, the number of nucleotides affected by PAPs could be hundreds of times higher than the number of SNPs for some pairs of isolates.

**Table 5.** The numbers of SNPs and PAPs unique to each of the isolates in the pair along with the total number of nucleotides in the PAPs

| Isolate 1/Isolate 2 | #SNP | #PAP | #Nucleotide<sup>a</sup> |
| --- | --- | --- | --- |
| FOC.TR4-5/FOC.TR4-1 | 69/85 | 65/379 | 21610/203003 |
| S1B8/JV14 | 202/153 | 109/68 | 33628/30997 |
| Leb1.2C/JV11 | 51/49 | 0/340 | 0/107249 |
| II-5/Vn-2 | 210/188 | 625/73 | 260086/42659 |
| Col2/Hainan.B2 | 211/254 | 555/24 | 268372/7094 |
| Col4/Col17 | 48/44 | 8/289 | 5022/114252 |
| My-1/La-2 | 121/68 | 88/0 | 40957/0 |
| Phi2.6C/Pak1.1A | 185/149 | 27/22 | 11269/8987 |

<sup>a</sup> The numbers before and after the slash (/) refer to the isolates before and after the slash in the pair, respectively.

Besides PAPs, we explored another source of structural variation – the copy number variation (CNV) of five common transposons in the isolate UK0001 and other 16 TR4 isolates. The five common transposons were the three elements mentioned previously, a SINE element and a LINE element. The representatives for these 5 elements in the UK0001 assembly were named: SINE of 650 bp at 3816788-3817437 of sequence accession VMNF01000006, LINE of 5893 bp at 3810881-3816773 of VMNF01000006, LTR1 of 8075 bp at 22,127-30,201 of VMNF01000011, and LTR2 of 3482 bp at 87797-91278 of VMNF01000013, and Helitron of 6122 bp at 66,262-72,383 of VMNF01000013, where LTR1 was a full-length version of the 5682-bp element mentioned previously. Note that SINE and LINE were 15 bp away.

The estimated copy numbers of the five transposons in the the isolate UK0001 and 16 other TR4 isolates are shown in Table 6. The copy number (found by Minimap2 in the UK0001 assembly) for SINE was 55, that for LINE was 14, that for LTR1 was 50, and that for Helitron was 32. The average sequencing coverage depth for the UK0001 dataset of short reads was less than 20, a cutoff value for ensuring accuracy in using read depth in copy number estimation (Wang et al., 2011); those for the rest were between 26 and 174. For each copy of each element, we also checked on the AT content of its surrounding region, which was defined to be centered in the copy and to be five times the length of the copy. For SINE, 46 of the 55 copies in the UK0001 assembly were in AT-neutral regions with AT contents of 45-52%; for Helitron, 25 of the 32 copies were in AT-neutral regions with AT contents of 47-52%; for LTR1, 42 of the 50 copies were in 53-71% AT-rich regions. LTR1 was significantly (E-value *< e*^*−*50^) similar to 44 AT-rich regions each with an AT content of at least 65%; the AT content of LTR1 was 53%. LTR1 was 78.4-81.9% identical (over 8,036-8,041 bp) to these three regions in Focb race 1 160527: 92,289-100,330 bp of race 1 contig 8, 55,507-63,543 bp of race 1 contig 8, and 3,574,210-3,582,252 bp of race 1 contig 9, which were geneless and 67-70% AT-rich. Table 6 indicates that the copy numbers of an element like SINE in some isolates of the same clade differed significantly.

**Table 6.** The copy numbers of five transposons in isolate UK0001 and 16 other TR4 isolates

| Isolate | SINE | LINE | LTR1 | LTR2 | Helitron |
| --- | --- | --- | --- | --- | --- |
| UK0001 | 57 | 13 | 53 | 10 | 28 |
| FOC.TR4-1 | 42 | 4 | 57 | 9 | 35 |
| FOC.TR4-5 | 13 | 1 | 41 | 7 | 18 |
| JV14 | 22 | 2 | 45 | 10 | 26 |
| JV11 | 18 | 3 | 47 | 11 | 28 |
| Leb1.2C | 26 | 2 | 47 | 11 | 28 |
| II-5 | 88 | 33 | 46 | 18 | 21 |
| Col2 | 12 | 7 | 25 | 10 | 17 |
| Col4 | 11 | 6 | 25 | 8 | 19 |
| Col17 | 12 | 8 | 25 | 12 | 14 |
| S1B8 | 60 | 21 | 32 | 11 | 9 |
| My-1 | 30 | 8 | 27 | 8 | 14 |
| Vn-2 | 6 | 1 | 32 | 9 | 13 |
| La-2 | 10 | 1 | 31 | 11 | 15 |
| Phi2.6C | 72 | 23 | 39 | 8 | 11 |
| Pak1.1A | 25 | 2 | 37 | 13 | 18 |
| Hainan.B2 | 6 | 4 | 33 | 11 | 10 |

In the Focb TR4 UK0001 assembly, we found that a region at 18,838-41,239 bp of supernumerary contig 13 was duplicated in reverse orientation at 2,598,351-2,620,752 bp of core contig 15. This perfect duplication was unique to isolate UK0001, not shared by any of the 16 other TR4 isolates. A large part of this duplicated region was also contained in a perfect match of 24,561 bp between a region at 14,135-38,695 bp of UK0001 supernumerary contig 13 and and a region at 2,760,458-2,785,018 of race 1 160527 supernumerary contig 2.

#### *F. oxysporum* f.sp *melonis*

In isolate Fom001 (van Dam et al., 2017), supernumerary contig 13 of 128.0 kb ends in a 5’ telomeric repeat. Contig 13 over positions 20.7 to 81.9 kb and over positions 81.8 to 111.8 kb displayed 99.99% identity to Forc016 chromosome RC over positions 0.1 to 61.3 kb and over positions 62.6 to 92.6 kb, respectively. The larger (with a length of 61 kb) of these two highly similar regions contained the 5’ subtelomere of chromosome RC (Fig. 1C), which was present at one or two ends of the 11 core chromosomes in isolate Forc016, but in a single copy (as an internal part of contig 13) in isolate Fom001. On the other hand, the 5’ subtelomere of contig 13 located at 1-19.3 kb, which was not present in isolate Forc016, was found at the telomere-containing ends of 17 contigs in isolate Fom001 at 98.69-99.76% identity for 6 of them over a length of 19 kb and for the rest over a length of 9.8-13.0 kb. Note that large portions of Forc016 chromosome RC were at 99.27-99.99% identity to parts of supernumerary contig 127 of 1,743,2 kb in isolate Fom001. Indeed, the mean SNP rates of windows in supernumerary contigs 13 and 127 were 26.63 and 20.19 times lower than that in the genome between isolates Fom001 and Forc016.

Within isolate Fom001, the 5’ subtelomere of contig 13 was 77% identical over positions 0.7 to 7.8 kb and 63% identical over positions 10.4 to 20.7 kb to the 5’ subtelomere of contig 22 over positions 1.2 to 8.1 kb and over positions 1.2 to 9.8 kb, where the 63%-identity match included a block of 1,483 bp at 100% identity. This revealed a tandem duplication in the 5’ subtelomere of contig 13, where both copies were similar to parts of the 5’ subtelomere of contig 22. Moreover, the 5’ subtelomere of contig 13 was 85% identical over one copy and 95% identical over the other copy to a region of supernumerary Forc016 contig 21 and a region of supernumerary Forc016 contig 52, respectively (Fig. 1C). Again within isolate Fom001, contig 22 of 1,267,8 kb was supernumerary; it was enriched for transposons and SM/pathogenicity genes. The 5’ 9.8-kb subtelomere of contig 22 was 99.27-99.96% identical to the 5’ subtelomeres of contigs 9, 15, 32, 86, 87, to a region of contig 107 from positions 9.9 to 19.7 kb, all in forward orientation, and to the 3’ subtelomere of contig 30 in reverse orientation.

Isolate Fom001 was more similar in core chromosome regions to the Fol lineage with isolate Fol007 than to other lineages including the Fom lineage (van Dam et al., 2016), and Fol D11 and Fom001 were 99.97% identical over a 20.5-kb region containing the conserved *EF-1α* gene. However, the top matches between supernumerary Fom001 contig 22 and the *Fusarium* genome assemblies were all but one with isolates Fom004, Fom005, Fom006, Fom011, Fom012, Fom013 and Fom016 at 99.83-99.99% identity over several regions of 31.9 to 42.7 kb. More specifically, the mean SNP rate (0.00037) of windows in the Fom001 genome between isolates Fom001 and Fol007 was 9.38 times lower than that (0.00347) between isolates Fom001 and Fom006. On the other hand, supernumerary Fom001 contigs 13, 22 and 127 were not present in isolate Fol007. Between isolates Fom001 and Fom006, the mean SNP rates of windows in those three contigs were 23.93, 2.73 and 3.49 times lower than that in the Fom001 genome. The same observation was made for isolate Fom005.

The 5’ subtelomere of Fom001 contig 13 was covered over part 1 (1 bp to 9.8 kb) by short reads from some of the other Fom isolates; the portion from 13 to 15 kb was not covered by short reads from any of them. The estimated copy number of part 1 was 34 in Fom005, 27 in Fom006, 0 in Fom009, 44 in Fom010, 0 in Fom011, 28 in Fom012, 24 in Fom013 and 28 in Fom016; there was a problem in mapping short reads with the Fom004 dataset. We found 1 to 3 SNPs in part 1 between Fom001 and each of the other 6 Fom isolates with multiple copies of part 1, and depths of low coverage from 1.3 to 1.6 kb for each isolate, suggesting variation over this region between Fom001 and the others. The estimated copy number of the 5’ subtelomere of Fom001 contig 22 was 0 in isolates Fom009, Fom010 and Fom011, and 4 to 7 in isolates Fom005, Fom006, Fom012, Fom013 and Fom016, with only 1 SNP at the same position between Fom001 and each of the other 5 isolates.

The 5’ 19-kb subtelomere of contig 13 contained 4 genes: two of them encoded for proteins with an MYND finger domain (4.1e-08 and 4.2e-10), and the other two encoded for hypothetical proteins of 1,340 and 1,255 residues. (In other words, each of the two copies in the 5’ subtelomere contained two genes, one encoding for an MYND finger domain and the other for a hypothetical protein.) About 8.7 kb downstream of the subtelomere was a gene (shared with Forc016 chromosome RC, see above) encoding for a 2,987-residue protein with DEAD (2.1e-28), Helicase C (2.3e-27), ResIII (2.8e-06) domains. The 5’ 9.8-kb subtelomere of contig 22 contained a gene encoding for a 403-residue protein with an MYND finger domain (1.3e-09) and another gene encoding for a hypothetical protein of 1,543 residues. About 4.5 kb downstream of the subtelomere was a gene encoding for a 1,317-residue protein with DEAD (1.2e-10), Helicase C (1.5e-07), and DUF3505 (6.2e-05) domains. Immediately downstream of this gene was another gene encoding for a 2,649-residue protein with DUF3505 (3.5e-13) and DEAD (2.5e-06) domains.

#### *F. oxysporum* f.sp. *cubense*

*F. oxysporum* f.sp. *cubense* (Focb) is a *forma specialis* of polyphyletic lineages causing Panama disease of banana (O’Donnell et al., 1998). One group of lineages is Focb race 1, which attacks a banana cultivar named ‘Gros Michel’ and caused the 20th century epidemic; another group is Focb tropical race 4 (TR4), which affects a banana cultivar named ‘Cavendish’ (resistant to race 1) and the hosts of race 1 (Zheng et al., 2018). Highquality genome assemblies were recently generated for Focb race 1 isolate 160527 (Asai et al., 2019) and for Focb TR4 isolate UK0001 (Warmington et al., 2019). Focb 160527 had a total of 15.7 Mb strong matches (of length above 1 kb and at identity above 98.7%) to isolate Forc016 (not in Focb), as compared to a total of 9.3 Mb strong matches to Focb TR4 UK0001, suggesting that Focb 160527 was more closely related to Forc016 than to Focb TR4 UK0001, which is consistent with the polyphyletic characterization of Focb. We observed a difference rate of 0.0167 between Focb race 1 160527 and TR4 UK0001 over a 20.5-kb region containing the conserved *EF-1α* gene.

The Focb race 1 160527 assembly consisted of 12 contigs with 20 terminal regions ending in telomeric repeats. One of the contigs, contig 2 of 5,885.8 kb with each terminal region ending in a telomeric repeat, was a supernumerary chromosome flanked on each side by reverse complementary subtelomeres of 8.5 kb at 99.98% complementarity. Note that this supernumerary chromosome was longer than all but one contig, with 7 of them assembled from telomere to telomere. The 3’ subtelomere of supernumerary contig 2 was at 98.83-99.98% identity to 20 contig terminal regions in forward or reverse orientation. Another contig, contig 12 of 1,261.1 kb, was also supernumerary with th same kind of reverse complementary subtelomeres as contig 2. The Focb TR4 UK0001 assembly was composed to 15 contig with 20 terminal regions ending in telomeric repeats. Contig 13 of 1,244.7 kb was supernumerary.

To see if part of the 3’ Focb 160527 supernumerary subtelomere was present in Focb TR4 isolates, short reads from three Focb TR4 isolates La-2, My-1 and Vn-2 (Zheng et al., 2018) were mapped onto the subtelomere as a reference. A 1.6-kb region of the subtelomere at a distance of 3.2 kb from its 3’ end was covered at a maximum read depth of 538 for La-2, 1,587 for My-1, and 1,988 for Vn-2. An estimated copy number for this region was 24 in La-2, 20 in My-1, 22 in Vn-2. A total of 14 SNPs between the reference and short reads, with each SNP shared by the three TR4 isolates, were found in this region. The Focb 160527 subtelomere contained a gene encoding for a 1454-residue protein with a DEAD domain (1.3e-08), a Helicase C domain (1.7e-08) and an OrsD domain (9.7e-05). The region of the subtelomere covered by short reads from the three TR4 isolates included all of the Helicase C domain and the C-terminal half of the DEAD domain. Although Focb 160527 and Focb TR4 UK0001 were not closely related, the three TR4 isolates contained a region in 20-24 copies that was at 99.1% identity to a region of the Focb 160527 subtelomere. The Focb 160527 assembly was used as a reference for mapping short reads from the three TR4 isolates, more than 64% percent of contig 2 (or 96% of contig 12) were not covered by short reads from each of the three TR4 isolates; about 18-58% percent of every other contig were not covered by short reads from each of the three TR4 isolates. Although Focb TR4 isolates had little variation with low SNP rates around 0.000005 (Zheng et al., 2018), two regions of Focb 160527 contig 2 were found with little variation with two of the three TR4 isolates but with significant variation with the other isolate. One was a 29.6-kb region starting at 3,645.9 kb of contig 2 that was covered with no SNPs at average read depths of 56.4 (for My-1) and 55.2 (for Vn-2) but at an average read depth of 0.1 for La-2. The other was a 47.4-kb region starting at 4,006.6 kb of contig 2 with one common SNP at average read depths of 18.3 (for La-2) and 54.8 (for Vn-2) but at an average read depth of 0.0 for My-1.

Besides transposons, supernumerary contig 2 contained genes encoding for 2 proteins each with a Cyclin N domain (6.4e-26 and 4.4e-07), 4 proteins each with a PARP/PARP reg domain (3.9e-32, 1.6e-20, 1.9e-20 and 2.7e-22), an HSP70 protein (1e-57), a protein with RCC1 and RCC1 2 domains (2.4e-66 and 5.6e-40), 3 proteins each with a Lactamase B domain 2.1e-06, 4.9e-11 and 6.4e-17), and 2 proteins each with a Lysm domain (3e-06 and 4.3e-15).

### HET domain genes

We collected data on the number, location and variation of HET domain genes in isolate Fol D11 to shed light on their origin and role. We predicted 135 HET domain genes with an e-value below 1.e-05 in the Fol D11 genome assembly. Of these genes, 23 (17%) were located in the supernumerary chromosomes or contigs, where a contig was classified as being supernumerary if its extent of coverage by short reads from isolates Fol069 and Fol072 was less than 60% and it was enriched for transposons and SM/pathogenicity genes. [At least 73% (or 74%) of each D11 core chromosome was covered by short reads from Fol069 (or Fol072), and 56.4% (or 55.1%) of supernumerary Fol D11 chromosome 14 was covered by short reads from Fol069 (or Fol072).] Of the remaining 112 HET domain genes, 91 were found in the 9 core chromosomes and 21 in core contigs 14 and 16.

We evaluated the variation between D11 and each of isolates Fol4287, Fol069, Fol072 and Fo47 by examining the coverage of the D11 genomic regions around each of the 91 HET domain gene loci by short reads from each of these 4 isolates. Here the genomic region around a gene was defined as an area centered in the gene and of a length twice that of the coding region. Except for a 3’ 500-kb end region of chromosome 1 with 3 HET domain genes and a 5’ 40-kb end region of chromosome 11 with 1 HET domain gene, the D11 genomic regions around the loci of the HET domain genes were covered at sufficient depths (*≥* 10) without any variation by short reads from Fol4287. However, some positions in each of the D11 genomic regions around 89 of the 91 HET domain genes were not covered by short reads from Fol069, Fol072 or Fo47. This shows that the core genomic regions around the HET domain genes tended to be free of variation between isolates D11 and Fol4287 from the same lineage, but were variable between D11 and each of isolates Fol069, Fol072 and Fo47 in different lineages from D11. The 91 HET domain genes were located in regions with SM/pathogenicity genes; for example, 40 of the 91 genes were located within 15 kb of a gene encoding for a major facilitator superfamily (MFS) or cytochrome P450 protein. Similar observations were made about the 21 HET domain genes in core contigs 14 and 16; for example, 16 of these genes were located within 15 kb of genes encoding for MFS or P450 proteins.

## Discussion

It has been shown through experiment that supernumerary chromosomes can be transferred between asexual lineages that are subjected to heterokaryosis and vegetative incompatibility (He et al., 1998; Akagi et al., 2009; Ma et al., 2010; Vlaardingerbroek et al., 2016a; van Dam et al., 2017). In another experimental study, strains with multiple copies of a supernumerary chromosome were produced by protoplast fusion and dosage effects of this chromosome in the fusion products were evident as increased virulence. The probability for the formation of multiple copies of a supernumerary chromosome by intrastrain anastomosis is expected to be higher than that by interstrain anastomosis, because vegetative incompatibility system inhibits heterokaryon formation between strains with different genetic backgrounds (Garmaroodi & Taga 2007). In this study, by sequence analysis we found instances of horizontal transfer for some regions of Focb TR4 supernumerary contig 13 between Focb race 1 and Focb TR4 isolates; those transfers provided novel genes in the formation of supernumerary contig 13 in the ancestor of the young Focb TR4 population. We characterized structural variation in the young asexual pathogen race of Focb TR4 with an extremely low rate of SNP: two or more structurally different copies of Focb TR4 supernumerary contig 13 in field isolates, transposon CNVs among isolates, and duplications of supernumerary chromosomal regions in core chromosomes. The existence of field isolates with multiple copies of supernumerary chromosomes in a young asexual pathogen race suggests that the horizontal transfer of supernumerary chromosomes by intrastrain anastomosis is ongoing and that such field isolates pose new challenges to disease management.

In several asexual populations of filamentous fungal pathogens with supernumerary chromosomes, supernumerary and core chromosomes are flanked on either side by highly similar reverse complementary long sequences. This subtelomeric sequence often contains the open reading frame of a helicase gene, like the Y’ element in the budding yeast, whose helicase is expressed many fold in the absence of telomerase (Yamada et al., 1998). Thus, this subtelomeric homology structure may have a role in maintaining telomeres for supernumerary chromosomes as well as core ones during vegetative reproduction. The structure also suggests homologous recombination as a mechanism for the exchange of genes between supernumerary chromosomes and the ends of core chromosomes; such an instance occurred in the Focb TR4 UK0001 isolate. A large number of HET domain genes preserved within isolates from the same population or *forma specialis* may have a role in marking their neighboring non-essential pathogenicity genes, because a loss of a DNA segment with a HET domain gene may result in vegetative incompatibility. In Fol, some isolates are variable around most HET domain genes but are more conserved in core and supernumerary subtelomeres as well as in whole supernumerary chromosomes. This suggests that the subtelomeres and supernumerary chromosomes are host-specific. Thus, the subtelomeric homology structure on the core and supernumerary chromosomes in several asexual species may be used as genomic signatures of host adaptation or host-switching.

In Fol and between Forc and Fom, a supernumerary chromosome or parts of it are more conserved than the core chromosomes. This, combined with the *in vitro* experimental evidence of HCT with the supernumerary chromosome (Ma et al., 2010 and van Dam et al., 2017), suggests that HCT of supernumerary chromosomes between different asexual lineages occurs in nature. These asexual lineages are formed to acquire and maintain supernumerary chromosomes through the duplication of subtelomeres at the ends of core chromosomes. The highly conserved subtelomeric structure also suggests that HCT occurs recently. The presence of a new subtelomere with a long insertion in some core and supernumerary chromosomes indicates that the subtelomeric homology structure evolves as an ongoing process to acquire and maintain the most-pathogenic supernumerary chromosome in the *forma specialis*. Consequently, HCT occurs frequently in asexual filamentous fungal pathogens with supernumerary chromosomes. Supernumerary chromosomes function as a powerful tool of evolution for non-essential but beneficial genes in filamentous fungi: they carry and evolve such genes across a wider species complex of filamentous fungi in a complementary mechanism to the Mendelian process. Some of these genes are homologs of essential genes, involved in DNA replication and repair, and gene expression and chromosome homeostasis. Other regions such as their reverse complementary subtelomeres come from supernumerary chromosomes in other lineages. Taken together, HET domain genes along with SM/pathogenicity genes are acquired by HCT of supernumerary chromosomes, and exchanged between core and supernumerary chromosomes by homologous recombination over subtelomeres.

Core chromosomes contained numerous G-C to A-T nucleotide changes indicative of RIP, while supernumerary chromosome did not, which is consistent with the study by Vanheule et al. (2016). No such changes were observed in the highly similar subtelomeric sequences of core chromosomes. An explanation consistent with these observations is given as follows. A filamentous fungal pathogen alternates between sexual and asexual cycles (Nieuwenhuis & James, 2016). During the sexual cycle, highly similar regions in its genome, like highly homologous subtelomeric regions, sustain mutations by processes like RIP. The pathogen evolves by genome wide homologous recombination within the population. During the asexual cycle, the pathogen first acquires (by HCT) supernumerary chromosomes with highly similar reverse complementary subtelomeric sequences, which are subsequently duplicated at its core chromosomes. Then the pathogen evolves by localized homologous recombination over subtelomeres and by HCT of supernumerary chromosomes within and between populations. The asexual cycle allows some homologs of essential genes as well as SM/pathogenicity genes to evolve through gene duplication and HCT, without being suppressed by processes like RIP. The asexual cycle could last for thousands of generations; it was estimated that the wild yeast *Saccharomyces paradoxus* had an asexual cycle of 1,000 generations (Tsai et al., 2008). Each asexual cycle corresponds with the emergence of a new asexual lineage. So the length of the asexual cycle may be related to the frequency of speciation events.

An explanation for the reverse complementary long sequences with a helicase gene at the ends of a supernumerary chromosome is that the helicase and long reverse complementary sequence structure are involved in maintaining the stability of the chromosome by lengthening its telomeres based on homologous recombination in vegetative cells. This arrangement is especially important when the supernumerary chromosome is transferred horizontally into a different population whose chromosome ends are not homologous to those of the supernumerary chromosome.

Certain asexual filamentous fungal pathogens adapt to their hosts by acquiring supernumerary pathogenicity chromosomes horizontally and duplicating the supernumerary subtelomeres at the ends of their core chromosomes. An evolution-guided strategy for controlling these pathogens is to develop a comprehensive identification method based on their host-specific subtelomere structures and to deploy the method globally to prevent them from disseminating their supernumerary pathogenicity chromosomes. Specifically, it is necessary to carefully handle plants whose fungal pathogens are found to possess supernumerary chromosomes that were the source of numerous instances of horizontal transfer in the past.

## Additional Information and Declarations

### Author Contributions

Xiaoqiu Huang conceived and designed the experiments, performed the experiments, analyzed the data, contributed reagents/materials/analysis tools, wrote the paper, prepared figures and/or tables, reviewed drafts of the paper.

### Data Availability

No new sequence data were generated in this project.

## Acknowledgments

This research was conducted during X.H.’s Faculty Professional Development Assignment in 2018-2019.

